# High CXCL6 drives matrix expression and correlate with markers of poor outcome in IPF

**DOI:** 10.1101/2021.06.22.449424

**Authors:** Harinath Bahudhanapati, Jiangning Tan, Rosa Marie Apel, Benjamin Seeliger, Xiaoyun Li, Ting-Yun Chen, Daniel Sullivan, John Sembrat, Mauricio Rojas, Tracy Tabib, Eleanor Valenzi, Robert Lafyatis, Chetan Jawale, Partha Biswas, John Tedrow, Taylor Adams, Naftali Kaminski, Wim A Wuyts, John F McDyer, Jonathan K Alder, Yingze Zhang, Mehdi Nouraie, Antje Prasse, Daniel J Kass

**Author notes:** To whom all correspondence should be addressed: Simmons Center for Interstitial Lung Disease. Division of Pulmonary, Allergy, and Critical Care Medicine. University of Pittsburgh School of Medicine, 200 Lothrop St, Pittsburgh.; E- mail. denotes equal contribution (Harinath Bahudhanapati, Jiangning Tan). denotes equal contribution (Antje Prasse, Daniel J Kass).

## Abstract

Signaling via G protein-coupled receptors (GPCRs) can modulate levels of cyclic adenosine monophosphate (cAMP) and shape the functions of fibroblasts in idiopathic pulmonary fibrosis (IPF). We have identified Chemokine (C-X-C) Motif Ligand 6 (CXCL6) as a potential pro-fibrotic GPCR ligand. We tested the function of CXCL6 in *ex vivo* human donor and fibrotic lung fibroblasts and in an animal model of pulmonary fibrosis. We also measured levels of CXCL6 in the blood and bronchoalveolar lavage (BAL) of patients with IPF. CXCL6 decreased cAMP levels in a dose-dependent manner in Donor and IPF Fibroblasts. CXCL6 mRNA and protein were localized to epithelial cells. Administration of mCXCL5 (LIX, murine CXCL6 homologue) to mice increased collagen synthesis with and without bleomycin. CXCL6 increased Collagen I and α-SMA levels in Donor and IPF Fibroblasts. Silencing of CXCR1/2 as well as Reparixin, a CXCR1/2 inhibitor, blocked effects of CXCL6. Treprostinil blocked effects of CXCL6 only on levels of α-SMA but not on Collagen I. CXCL6 levels in the BAL of two separate cohorts of patients with IPF was associated with poor survival. We conclude that high CXCL6 drives fibroblast function and correlates with poor outcomes in IPF.

## INTRODUCTION

Idiopathic pulmonary fibrosis (IPF) is a progressive and devastating disorder characterized by the unremitting accumulation of activated fibroblasts in the lung leading to scarring and ultimately respiratory failure. One therapeutic approach has been to target activation of the fibroblast compartment through G protein-coupled receptor signaling. Specifically, agents that couple to Gαs and increase cyclic adenosine monophosphate (cAMP), such as prostaglandin E2 (PGE2) and dopamine, have been considered to be potential anti-fibrotic agents that decrease collagen synthetic activity (1–3). We and others have had a long-standing interest in the anti-fibrotic potential of the hormone relaxin, which is also a Gαs agonist via its receptor RXFP1. But despite the strength of the pre-clinical evidence supporting the anti-fibrotic potential of agonists of Gαs, relaxin failed in a clinical trial for the fibrotic disorder, systemic sclerosis (SSc) (4). One potential explanation for this observation is that relaxin/RXFP1 signaling, like PGE2 signaling, is attenuated in pulmonary fibrosis (2, 5–9). Another potential explanation is that the relative reduction of Gαs signaling may presage a shift in GPCR tone to other G proteins, such as Gαi, for example, that have been associated with reduced levels of cAMP and thus “pro-fibrotic” signaling (10). This leads us to postulate that there may be a “second hit,” perhaps through Gαi, which antagonizes Gαs pathways, to synergize with reduced RXFP1 signaling and promote fibrosis. Because there has been considerably less focus on *positive* regulators of pro-fibrotic signals through Gαi-coupled receptors, we wondered if there is another, and possibly more potent, pro-fibrotic Gαi signal to augment the reduction of cAMP in fibrosis?

We previously published that IPF patients (n=134) with the lowest *RXFP1* gene expression exhibited the most impaired pulmonary function and were predicted to experience the highest mortality rate (8, 9). We reasoned that these “*RXFP1*-low” patients would be characterized by a distinct “pro-fibrotic” gene expression profile that may include dysregulation of several profibrotic genes, including genes encoding soluble signals that would augment the loss of cAMP in the mesenchymal compartment. We identified C-X-C Motif Chemokine Ligand 6 (*CXCL6*) (9), a member of the IL-8 family of chemokines, to be one of the most highly expressed genes in *RXFP1-*low patients. Because *CXCL6* codes for a chemokine which signals via the Gαi-coupled receptors CXCR1/2 (11), we tested the hypothesis that dysregulation of CXCL6 signaling via CXCR1/2 would promote fibroblast activation and correlate with markers of worse clinical outcomes in IPF.

## RESULTS

### CXCL6 expression is increased at the level of mRNA and protein in IPF and decreases fibroblast cAMP levels

In this first experiment, we tested the hypothesis that baseline levels of cAMP were lower in IPF fibroblasts than in Donor fibroblasts. Cells were isolated as described in the Materials and Methods section and cultured in complete serum. When compared to Donor lung Fibroblasts, IPF lung fibroblasts showed significantly lower cAMP levels (**Figure 1A**). We identified C-X-C Motif Chemokine Ligand 6 (*CXCL6*) (9), a member of the IL-8 family of chemokines, to be one of the most highly expressed genes in *RXFP1-*low patients. Because the receptors for CXCL6 couple to Gαi and are predicted to reduce cAMP, we prioritized CXCL6 for further testing. Based on the Lung Genomics Research Consortium microarray data (12). *CXCL6* gene expression is significantly increased in IPF compared to Donor lungs **(Figure 1B)**. In comparison to *CXCL6* from the LGRC dataset, other members of the IL-8 family including *CXCL5* and *PPBP* are mildly increased, but *CXCL2 and CXCL3* show reduced expression. *CXCL8,* which codes for IL-8, is not differentially expressed in IPF at the mRNA level **(Figure 1C)**. Next, we determined if CXCL6 is increased at the protein level. We generated whole lung protein lysates of IPF and Donor lungs and subjected these lysates to immunoblotting with an antibody that recognizes CXCL6 and CXCL5. By immunoblot, we found increased expression of CXCL5/6 in IPF lungs **(Figure 1D, E**). Next, we determined if there is increased protein expression of CXCL5, the mouse homologue of human CXCL6 (13). By BlastP, mCXCL5, is 64% identical to hCXCL6 and represents the closest match (*data not shown*). By immunoblot of lysates of bleomycin-injured lungs, we observed a significant increase in mCXCL5 levels compared to uninjured **(Figure 1F, G)**. Because the CXCL6 receptors are known to couple to the inhibitory G protein, Gα_i_ (11), we next asked if stimulation of lung fibroblasts with CXCL6 would decrease cAMP. Donor and IPF lung Fibroblasts were stimulated with CXCL6 at concentrations that range from 1 nM to 1 μM, and cells were processed to detect cAMP. Stimulation of Donor or IPF lung Fibroblasts with CXCL6 lead to a decrease in intracellular cAMP in a dose-dependent and saturable manner with IC_50_ of 1.7 nM for Donor (n=6) and 82 nM for IPF (n=3) **(Figure 1H and I).** Taken together, these data support the hypothesis that CXCL6 is up-regulated in IPF and may function to drive down fibroblast levels of cAMP.

**FIGURE 1.**
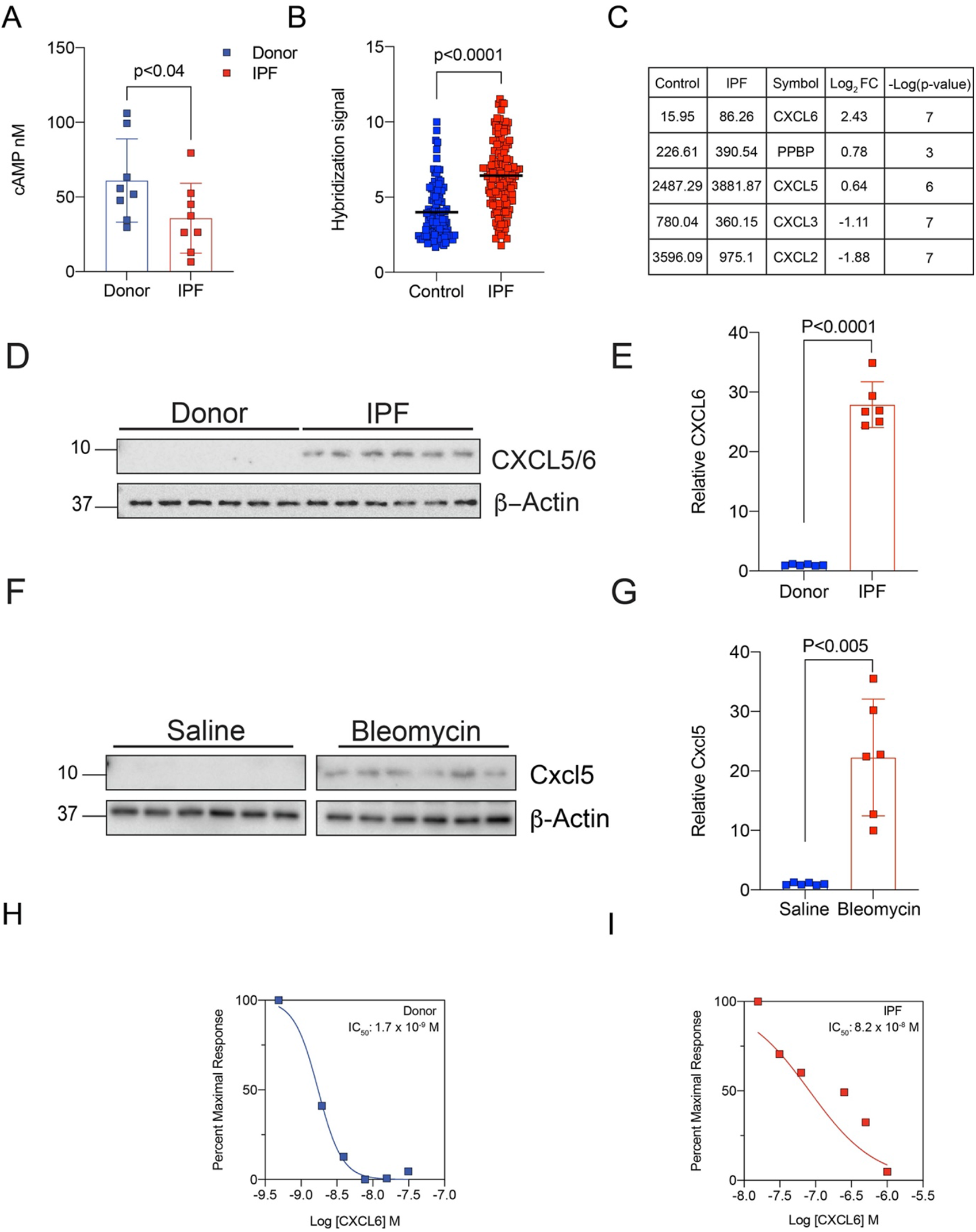
CXCL6 decreases cAMP levels in human lung fibroblasts. (A) In unstimulated cells, cAMP levels are significantly lower in IPF lung fibroblasts compared to Donor lung fibroblasts (Bootstrapped (resampling); p<0.04; N=8) (B) LGRC data shows that *CXCL6* expression is elevated in the IPF patients (N=134) compared to normal controls (N=108) (P<0.0001, Mann-Whitney). (C) LGRC data table showing differentially expressed members of the IL-8 family of chemokines. (D) Lysates of whole lung from human donors and IPF patients (N=6) were probed with an antibody that detects CXCL5 (13kDa) and CXCL6 (12kDa). (E) Densitometry of CXCL6 from (*D*) Data are expressed as mean ± SD., normalized to β-actin. Whole lung tissue extracts were immunoblotted for CXCL6 expression. CXCL6 is significantly highly expressed in lungs of IPF patients compared to Donor lungs (Mann Whitney test; p<0.0001, N=6). (F) Lysates of whole lung from mice that received Saline or Bleomycin were probed with an antibody that detects CXCL5/6. Bleomycin injury induced expression of CXCL5. (G) Densitometry of CXCL5/6 from (*F*) Data are expressed as mean ± SD., normalized to β-actin. Animals were injured with bleomycin or saline controls on day 0. CXCL5 is expressed following bleomycin injury compared to saline controls (Mann-Whitney test; p<0.005, N=6). (H) Donor lung fibroblasts were stimulated with CXCL6 for 20min and then lysed for determination of cAMP levels. CXCL6 decreased cAMP levels in a dose dependent manner. IC_50_ values were calculated for Donor fibroblasts and reported on the semi-log graph (N=6; Blue squares; mean IC_50_= 1.7 nM). (I) IPF lung fibroblasts were stimulated with CXCL6 for 20min and then lysed for determination of cAMP levels. CXCL6 decreased cAMP levels in a dose dependent manner. IC_50_ values were calculated for IPF fibroblasts and reported on the semi-log graph (N=3; Red squares; mean IC_50_= 81.9 nM).

### Single cell (sc) RNASeq demonstrates that CXCL6 mRNA is expressed in epithelial cells

Previously reported scRNAseq data of IPF(14) (n=32) and control (n=28) lung cells were downloaded from GEO (Accession: GSE136831) and re-analyzed to investigate expression patterns of CXCL6 (**Figure 2A-B**). Among the cell types with the highest CXCL6 expression across both control and IPF cells were goblet cells (normalized expression of 0.596, detected in 42% of cells), mesothelial cells (normalized expression of 0.269; in 27.6% of cells), disease-associated aberrant basaloid cells (normalized expression of 0.213, 27% of cells) and club cells (normalized expression of 0.142; in 12.5% of cells) (**Figure 2C**; Supplement: **CXCL6_CellTypeExpressionSummaryStats.txt**). Differential expression analysis in each cell type between IPF and control revealed that CXCL6 is significantly upregulated at both the cell and subject-level of IPF goblet (cell p-value=0.0193, subject p-value: 0.0047) and club cells (cell p-value: 0.00024, subject p-value: 0.00083). Fold change differences were notably stronger among IPF goblet cells (cell logFC: 0.46; subject logFC: 0.24) when compared to club cells (cell logFC: 0.18; subject logFC: 0.078) (**Figure 2D**; Supplement: **IPFvsCtrl_scRNAseq_CXCR6_WilcoxRankSum.txt**).

**FIGURE 2.**
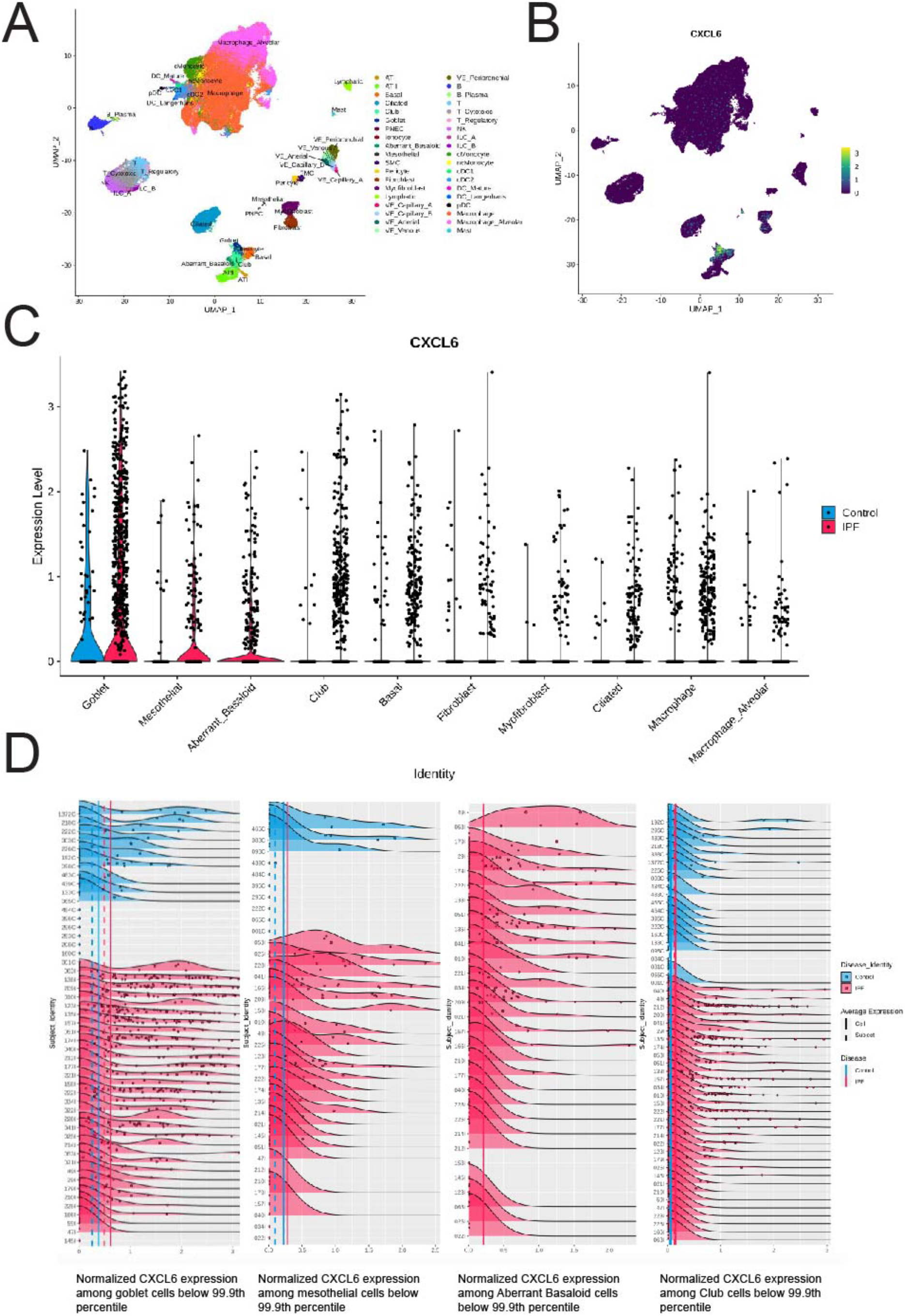
CXCL6 is expressed predominately in lung epithelial cells. (A) Uniform Manifold Approximation and Projection (UMAP) of 239,707 cells from 28 control and 32 IPF lung samples, with cells labeled as one of 38 previously described cell types. (B) UMAP of all cells, labeled by normalized CXCL6 expression. (C) Violin plots representing the distributions of normalized CXCL6 gene expression among the highest expressing cell types, split by control or IPF. Cell types are ranked in descending order of average CXCL6 expression. (D) Density plots representing the distribution of normalized CXCL6 expression across the top ranking cell types (left to right: goblet, mesothelial, aberrant basaloid and club) across control and IPF lung samples. Samples are grouped by disease then arranged in descending order of average expression. Solid and dashed lines correspond to cell and subject averages respectively, for each disease group.

To further corroborate the localization of *CXCL5/6* expression in IPF lungs, we also queried our recently published IPF scRNASeq dataset (**Figure 3A**) (15). *CXCL6* was expressed predominately in goblet cells, ciliated cells, basal cells, club cells, type I cells, and fibroblasts (16). For comparison, we also analyzed *CXCL5* gene expression by scRNASeq and found that *CXCL5* is expressed in inflammatory cells including macrophages (**Figure 3B**). By immunohistochemistry with an antibody that recognizes both CXCL5 and CXCL6, we observed staining in epithelial cells and macrophages **(Figure 3C)**. Immunostaining was observed in epithelial cells that overly fibroblastic foci which corroborates the scRNASeq data showing expression in disease-associated aberrant basaloid cells. We suggest that the staining that we observed in macrophages may represent CXCL5 and that the staining of epithelial cells likely represents CXCL6. Taken together, CXCL6 expression is predominately localized to epithelial cells in IPF, including the metaplastic epithelial cells adjacent to fibroblastic foci.

**FIGURE 3.**
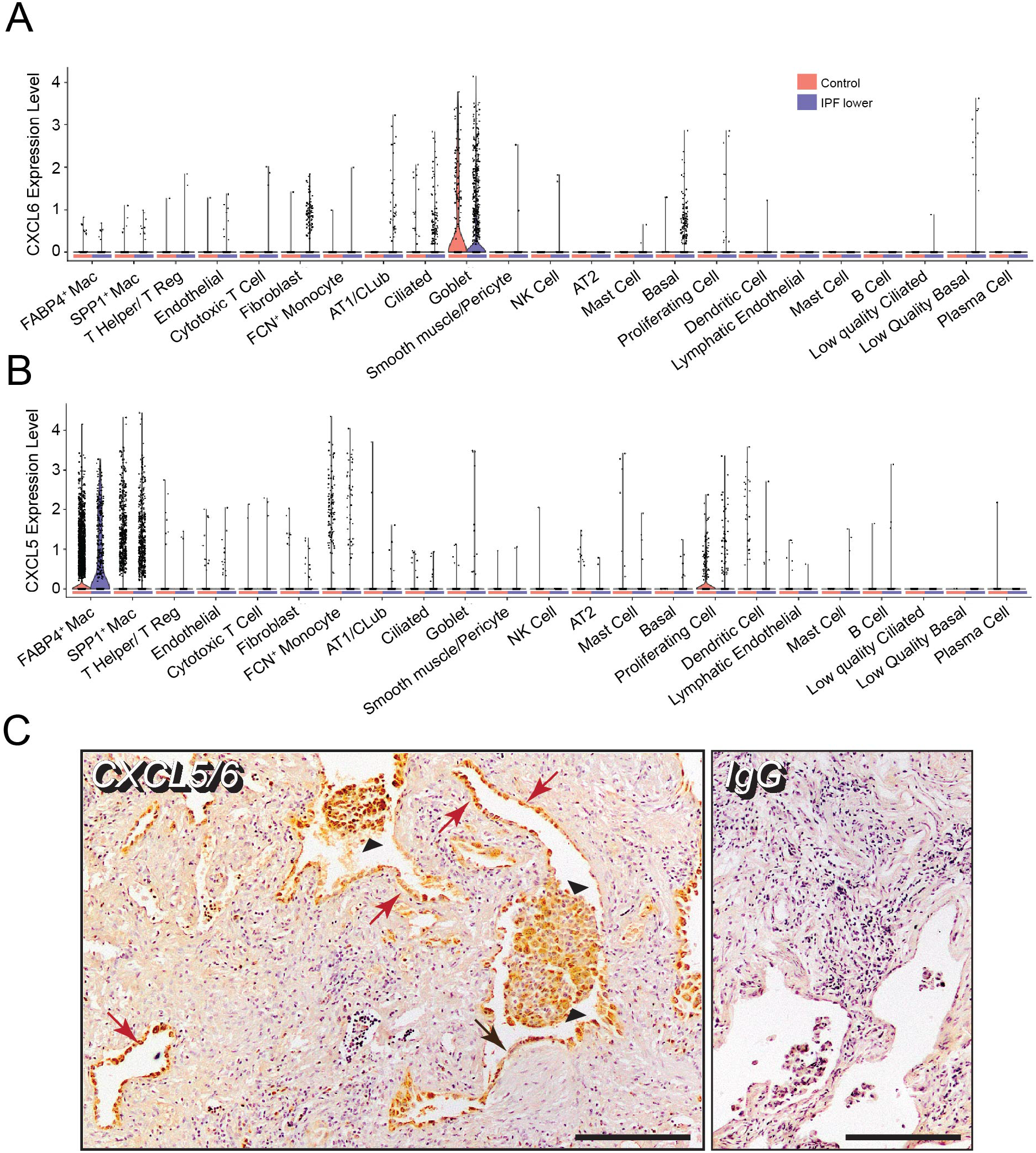
CXCL6 is expressed in lung epithelial cells while CXCL5 is expressed in macrophages. (A) By single cell RNASeq, *CXCL6* mRNA is present in goblet cells, ciliated cells, basal cells, club cells, type I cells, and fibroblasts (split violin plot) (B) By single cell RNASeq, *CXCL5* mRNA is present in inflammatory cells and macrophages (split violin plot) (C) Representative immunohistochemistry (IHC) for CXCL6 in IPF lungs shows staining in epithelial cells (red arrows), in epithelial cells adjacent to a fibroblastic focus (black arrows), and in macrophages (black arrowheads). (Right) Non-immune IgG shows an absence of staining (inset bar=200μm, magnification x100).

### Administration of mCXCL5 to mice increases collagen synthesis with and without bleomycin

Our immunohistochemistry suggested that CXCL6 from epithelial cells may stimulate the fibroblasts in fibroblastic foci. We have also shown that CXCL6 reduces fibroblast cAMP indicating that these cells are responsive. In this next experiment, we determined if murine CXCL5, the mouse homologue of human CXCL6, would increase collagen synthetic activity *in vivo* (*2*). Male and female mice were exposed to bleomycin or saline control by transnasal aspiration as previously published (9, 17, 18). On Days 7, 10, and 13 after injury, mCXCL5, or Saline control, was administered also by transnasal aspiration (*Schematic***; Figure 4A**). On day 14, the animals were euthanized, and the lungs were collected for histology, determination of collagen content, and quantification of inflammatory cells. By histology, we did not observe clear differences between uninjured animals treated with mCXCL5 or the vehicle control **(Figure 4B, C)**. Patchy injury was observed in bleomycin-injured mice treated with the vehicle control **(Figure 4D, E)**. In bleomycin-injured animals treated with mCXCL5, we observed more confluent areas of injury **(Figure 4F, G)** and the accumulation of inflammatory cells including neutrophils (**Figure 4H**). Next, we determined acid-soluble (new) collagen content in these lungs. We observed that uninjured animals treated with mCXCL5 exhibited similar collagen levels to bleomycin-injured animals treated with the vehicle control (**Figure 4I**). Animals injured with bleomycin and treated with mCXCL5 were characterized by the highest levels of collagen content. Next we analyzed the inflammatory cell content by flow cytometry of the BAL **(Figure 4J-M)**. We have consistently found that neutrophil (Gr1+) numbers are very low at 14d after bleomycin injury (9, 17, 18) **(Figure 4J)**. In the absence of bleomycin, mCXCL5 increased neutrophil numbers but not to a significant extent. However, in addition to bleomycin, mCXCL5 was associated with a significant accumulation of neutrophils. We also observed increased numbers of CD68+ **(Figure 4K)**, CD3+ **(Figure 4L)**, and B220+ **(Figure 4M)** cells in bleomycin-injured and mCXCL5-treated mice. These data show that mCXCL5 increases collagen synthetic activity in the lung. In bleomycin-injured animals, mCXCL5 also increased inflammatory cell numbers.

**FIGURE 4.**
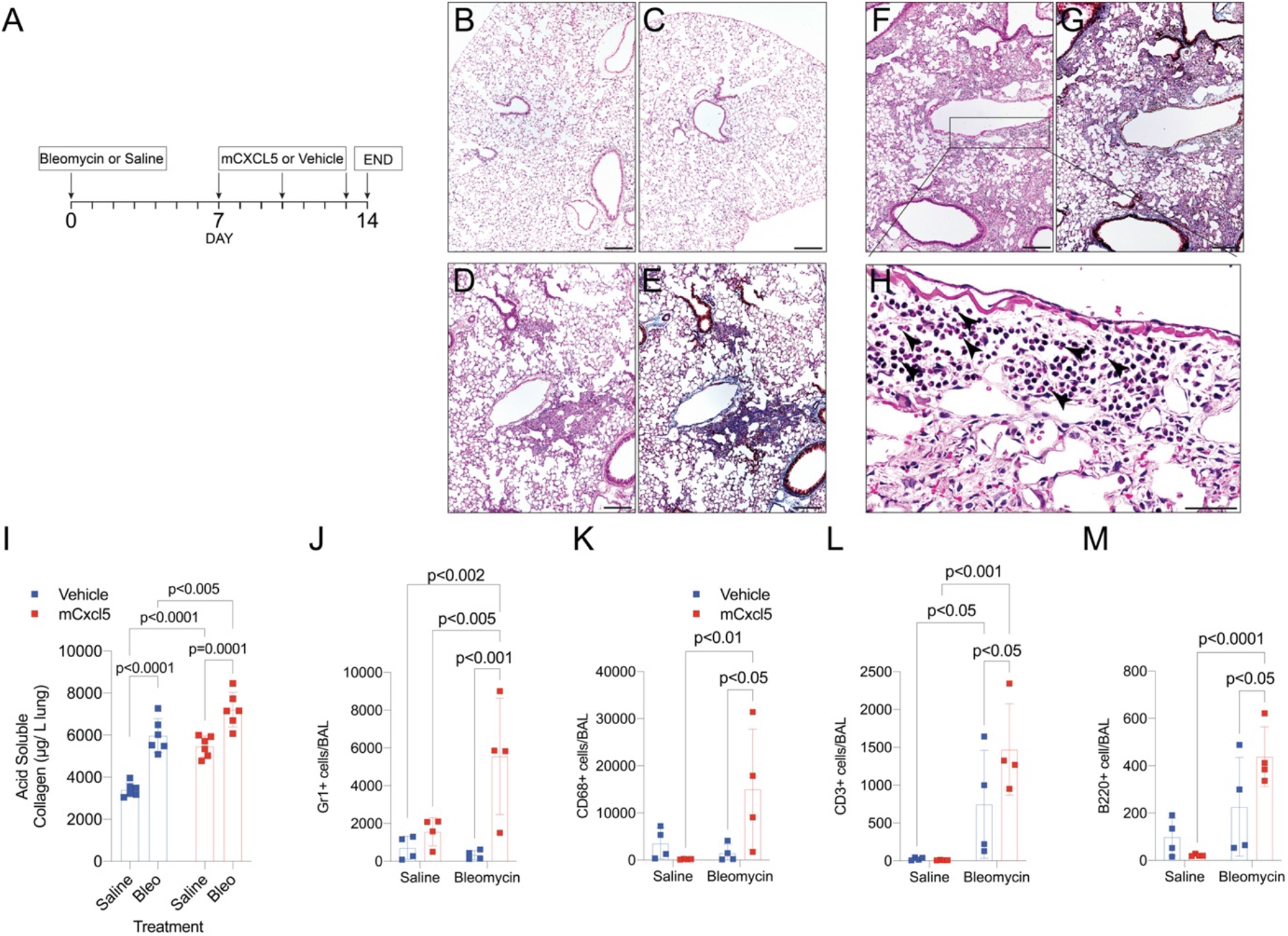
Mouse CXCL5, the homologue of human CXCL6, increases lung collagen in mice. (A) Schematic of the experiment. H&E staining of lungs at 14 d after bleomycin injury in WT animals (black inset scale bar, 200 μm; original magnification X100; upper panels) (B) Saline + Vehicle or (C) Saline + murine CXCL5. (D) H&E and (E) trichrome images of mice injured with bleomycin and treated with a vehicle control. (F) H&E and (G) trichrome images of mice injured with bleomycin and treated with murine CXCL5. (H) Outset image from F (black inset scale bar, 50 um; original magnification X400). (I) Acid-soluble collagen content of lungs from saline- or bleomycin-injured treated with vehicle or mCXCL5. P-values are indicated by two-ANOVA and followed by Fisher’s LSD post-hoc testing. (J) Flow cytometry: BAL was collected at day 14 for flow cytometry to detect (J) neutrophils (Gr1+), (K) macrophages (CD68+), (L) T-cells (CD3+), and (M) B-Cells (B220+), (N=4/group, by two way ANOVA followed by Fisher’s LSD post-hoc testing).

### CXCL6 and TGFβ increases Collagen I and α-SMA in Donor and IPF lung fibroblasts

To determine if lung fibroblasts respond to CXCL6, we stimulated Donor (n=3) and IPF lung fibroblasts (n=3) with CXCL6 (50ng/mL) with and without TGFβ (2ng/mL) **(Figure 5A)**. Lysates were subjected to immunoblotting (IB) for Collagen I and PPIA. Stimulation with CXCL6 alone increased Collagen I in both Donor and IPF Fbs. Incubation with CXCL6 and TGFβ, alone and cooperatively, increased collagen I expression (**Figure 5B**). CXCL6 treatment showed a significant increase in expression of Collagen I protein in Donor Fbs and IPF lung Fbs (p<0.01, n=3). A combination CXCL6 with TGFβ increased Collagen expression significantly in both Donor and IPF compared to CXCL6 treatment alone. Densitometry of α-SMA **(Figure 5*C*)** showed that CXCL6 treatment significantly increased expression of α-SMA protein in Donor lung Fbs while a combination CXCL6 with TGFβ resulted in a significant increase in α-SMA expression in Donor Fbs. These data show that CXCL6 increases fibroblast expression of collagen and α-SMA and is additive to the effects of TGFβ.

**FIGURE 5.**
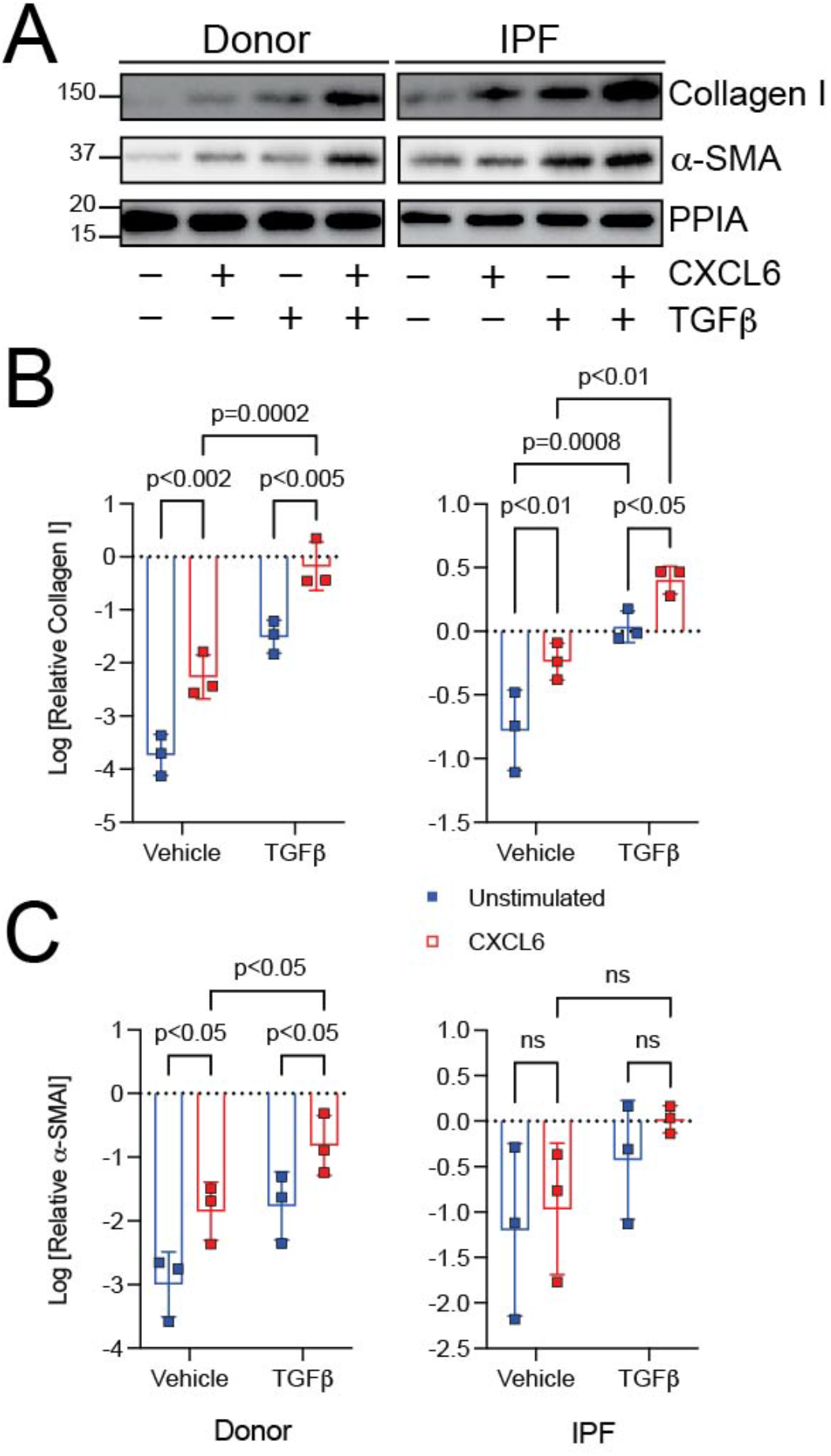
CXCL6 and TGFβ increases Collagen I in Donor and IPF lung fibroblasts. (A) Donor and IPF lung Fbs were serum starved and incubated in the presence of CXCL6 and/or TGFβ. Cells were processed for immunoblotting (IB) for Collagen I and PPIA (loading control). CXCL6 and TGFβ stimulation increase collagen levels to a greater extent than either alone. (B) Densitometry of Collagen I (*B*) from (*A*) data are log-transformed and expressed as mean ± SD., normalized to PPIA. CXCL6 increased expression of Collagen I protein in Donor (p<0.02) and IPF (p<0.01). A combination CXCL6 with TGFβ increased Collagen expression significantly in both Donor (p=0.0002, n=3) and IPF (p<0.01, n=3) compared to CXCL6 treatment alone (data analyzed by two-way ANOVA for Collagen I, followed by Fisher’s LSD post-hoc test). (C) Densitometry of α-SMA (*C*) from (*A*) data are log-transformed and expressed as mean ± SD., normalized to PPIA. CXCL6 treatment slightly increased expression of α-SMA protein in Donor. A combination CXCL6 with TGFβ increased α-SMA expression significantly in Donor (p<0.05, n=3) compared to CXCL6 treatment alone (data analyzed by two-way ANOVA for α-SMA, followed by Fisher’s LSD post-hoc test).

### CXCL6 increases expression of Collagen I and α-SMA in Donor and IPF fibroblasts via CXCR1/CXCR2

Myofibroblast activation (19–26), ECM accumulation (27–29) and contractility (30–32) are central to IPF pathogenesis. To determine if human lung fibroblasts respond to CXCL6, we stimulated Donor and IPF lung fibroblasts with CXCL6 (50 ng/mL or 6.2 nM). This dose was chosen based on our observed IC_50_ data and because CXCL6 has been shown to activate and internalize CXCR1/2 at higher doses (33). However, 50-100 ng/ml range has been established as an activating dose that did not affect internalization (34). In this experiment, we also determined if any CXCL6-induced effects on fibroblast function were dependent on expression of the CXCL6 receptors CXCR1 and CXCR2. Cells were transfected with non-targeting siRNA oligonucleotides or CXCR1 or CXCR2-targeting siRNA complexes. After 48h, lysates were subjected to IB for Collagen I, α-SMA, CXCR1/2 and PPIA **(Figure 6A-C and Supplemental Figure S1)**. Stimulation of both Donor and IPF fibroblasts with CXCL6 alone increased Collagen I. In Donor fibroblasts, the effect of CXCL6 on Collagen I was diminished by silencing of CXCR2. In IPF fibroblasts, CXCL6-induced increases in collagen expression were blocked by silencing of both CXCR1 and CXCR2. CXCL6 significantly increased α-smooth muscle actin (α-SMA) expression in Donor Fibroblasts. In IPF Fibroblasts, CXCL6 was associated with a borderline increase in α-SMA. Silencing of CXCR1 and 2 was associated with less CXCL6-induced expression of α-SMA across both Donor and IPF fibroblasts. Taken together, these data indicate that stimulation of fibroblasts is associated with increased expression of collagen and α-SMA and that these CXCL6-induced effects are dependent on expression of CXCR1 and CXCR2.

**FIGURE 6.**
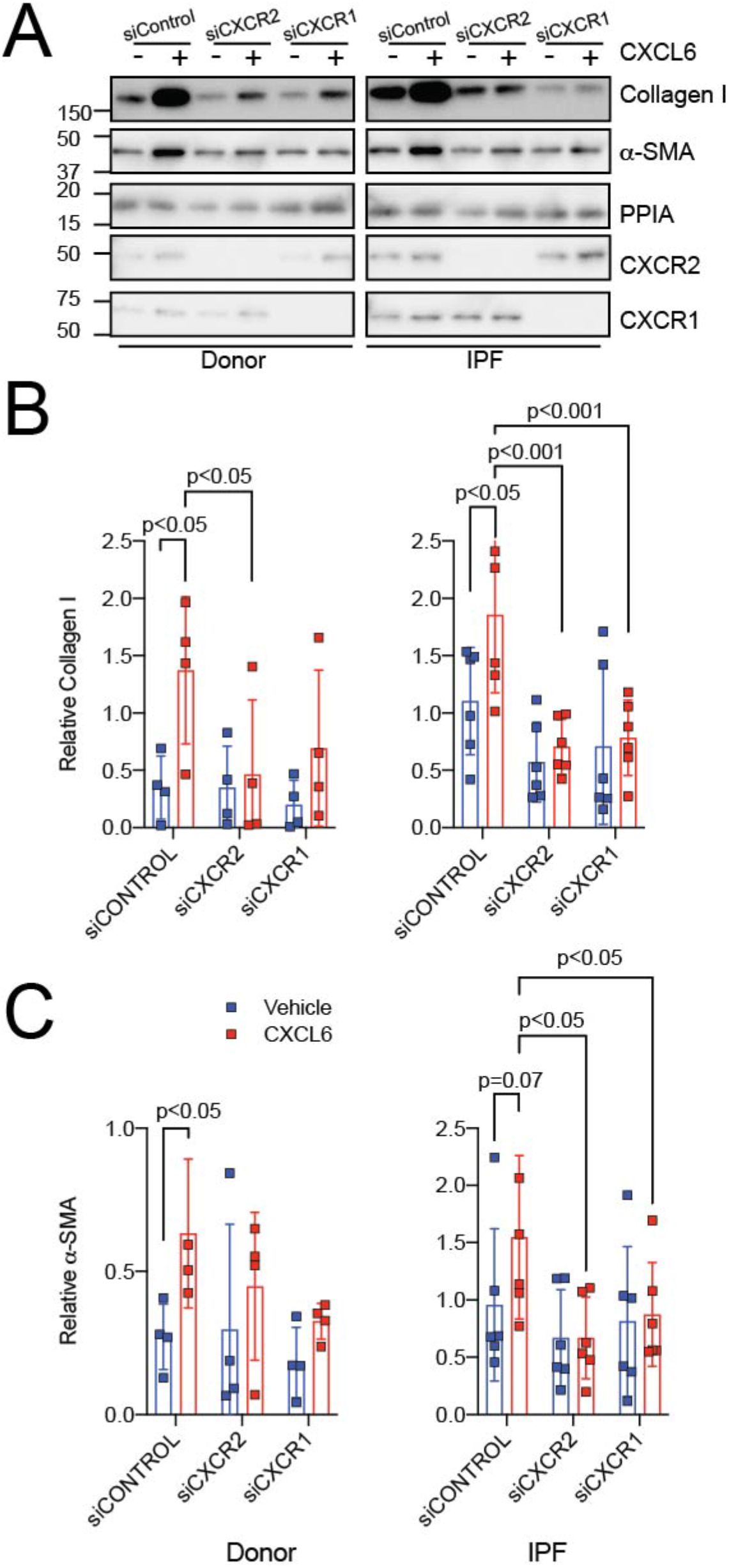
CXCL6 increases Collagen I and α-SMA in Donor and IPF fibroblasts, which can be blocked by silencing of CXCR1/CXCR2. (A) Donor and IPF lung Fibroblasts were transfected with CXCR1 or CXCR2-targeting siRNA or scrambled controls, serum starved, and incubated in the presence of CXCL6 (50 ng/mL, *red*) or a vehicle control (*blue*). Cells were processed for immunoblotting (IB) for Collagen I, α-smooth muscle actin (α-SMA), CXCR1, CXCR2, and house-keeping PPIA. CXCL6 increased collagen and α-SMA levels in Donor and IPF Fibroblasts. Silencing of CXCR1/2 decreased CXCL6-induced increases in collagen and α-SMA. Representative blot of experiments with 6 independent IPF and 4 independent Donor Fibroblasts. (B) Densitometry of Collagen I from Donor (*left*) or IPF (*right*) fibroblasts normalized to PPIA from (*A*) expressed as mean ± SD. Post-hoc comparisons are noted in the figure. Data were analyzed by two-way ANOVA, followed by Fisher’s LSD post-hoc test. (C) Densitometry of α-SMA from Donor (*left*) or IPF (*right*) fibroblasts normalized to PPIA from (*A*) expressed as mean ± SD. Post-hoc comparisons are noted in the figure. Data were analyzed by two-way ANOVA, followed by Fisher’s LSD post-hoc test.

### Reparixin blocks effects of CXCL6

Reparixin, a noncompetitive allosteric inhibitor of IL8, is more potent in inhibiting CXCR1 than CXCR2 (35). Here, we determined if chemical blockade of CXCR1/2 would phenocopy our observations with siCXCR1/2. We incubated Donor and IPF Fibroblasts in the presence or absence of CXCL6 with or without the CXCR1/2 blocker reparixin (1μM) (36) **(Figure 7A**). In Donor and IPF fibroblasts, CXCL6 increased expression of collagen I **(Figure 7B**) and α-SMA **(Figure 7C**). Incubation of cells with Reparixin decreased CXCL6-induced increases in collagen I and α-SMA. These data suggest that CXCL6 increases matrix and α-SMA expression in both Donor and IPF fibroblasts and that the effects of CXCL6 can be blocked by reparixin.

**FIGURE 7.**
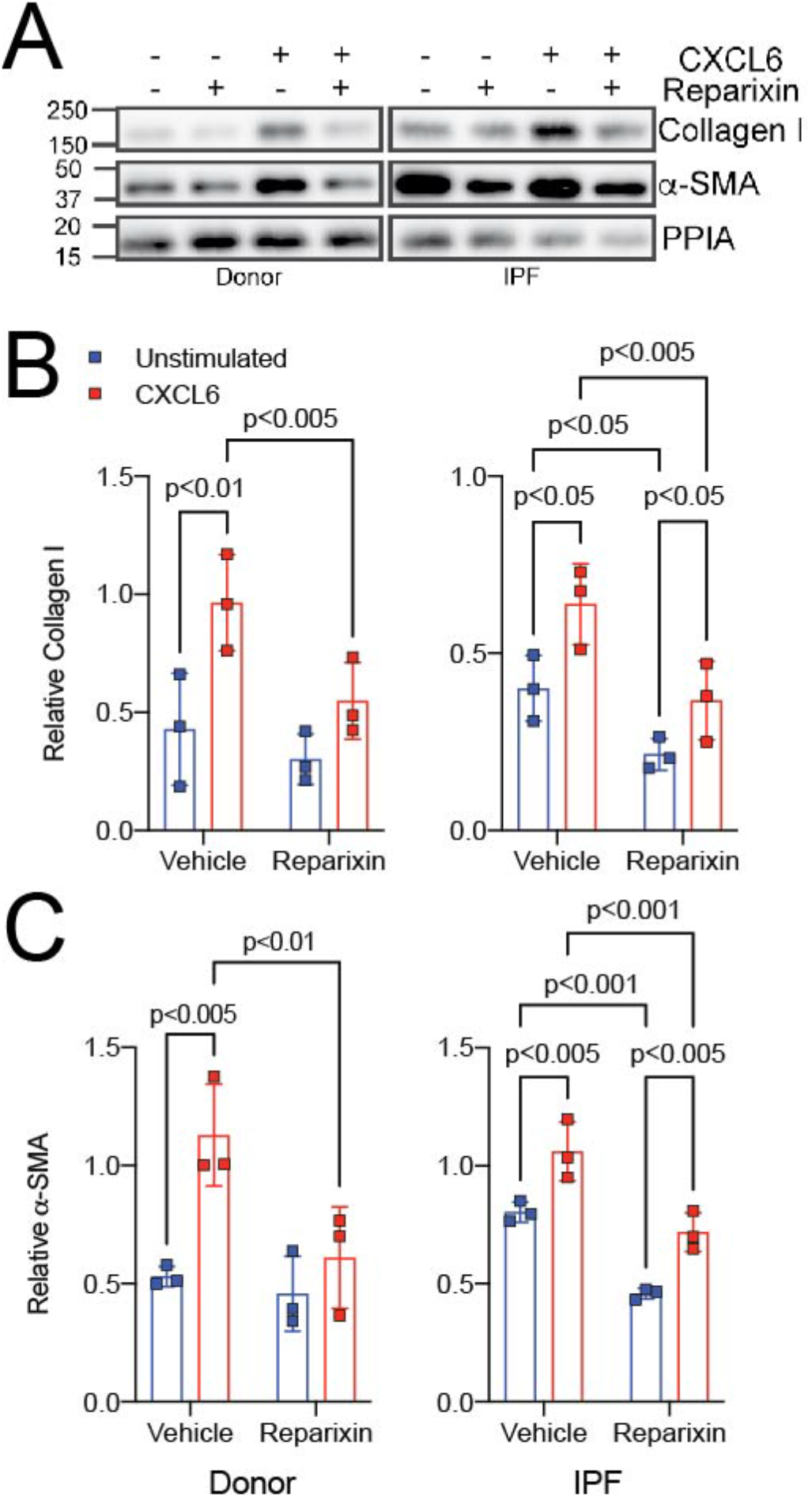
Reparixin reverses CXCL6-dependent increase in Collagen I in Donor and IPF fibroblasts. (A) Donor and IPF lung Fibroblasts were serum starved, stimulated with CXCL6 in the presence of the CXCR1/2 inhibitor, reparixin. Reparixin blocked CXCL6-induced effects on collagen and α-SMA. Representative blot of experiments with 6 independent IPF/ 4 Donor Fibroblasts. Cells were processed for immunoblotting (IB) for Collagen I, α-SMA and cyclophilin A. CXCL6 and TGFβ stimulation increase collagen levels to a greater extent than either alone. (B) Densitometry of Collagen I from Donor (*left*) or IPF (*right*) fibroblasts normalized to PPIA from (*A*) expressed as mean ± SD. Post-hoc comparisons are noted in the figure. Data were analyzed by two-way ANOVA, followed by Fisher’s LSD post-hoc test. (C) Densitometry of α-SMA from Donor (*left*) or IPF (*right*) fibroblasts normalized to PPIA from (A) expressed as mean ± SD. Post-hoc comparisons are noted in the figure. Data were analyzed by two-way ANOVA, followed by Fisher’s LSD post-hoc test

### Treprostinil blocks CXCL6-induced increases in α-SMA but not collagen I

If decreasing cAMP is associated with pro-fibrotic functions in fibroblasts, we reasoned that incubation of lung fibroblasts with cAMP would then decrease CXCL6-induced expression of collagen and α-SMA. To determine the effects of cAMP on CXCL6-induced increases in α-SMA and collagen I in lung fibroblasts, IPF and Donor Fbs were stimulated with CXCL6 in the presence of treprostinil, the synthetic analogue of the Gαs/cAMP agonist prostacyclin (PGI2) in order to block the effect of CXCL6 (**Figure 8A**). The prostacyclin receptor is expressed in lung fibroblasts (37, 38). Cells were pretreated either with vehicle or 10 mM of treprostinil (Sigma) prior to incubation with vehicle or recombinant human CXCL6 (50 ng/ml). Cells were lysed after 48 hrs for determination of α-SMA and collagen by immunoblotting. Treprostinil blocked CXCL6-induced increases in α-SMA but not in collagen I. These data indicate that CXCL6 increases collagen I expression independently of cAMP (**Figure 8B-C**).

**FIGURE 8.**
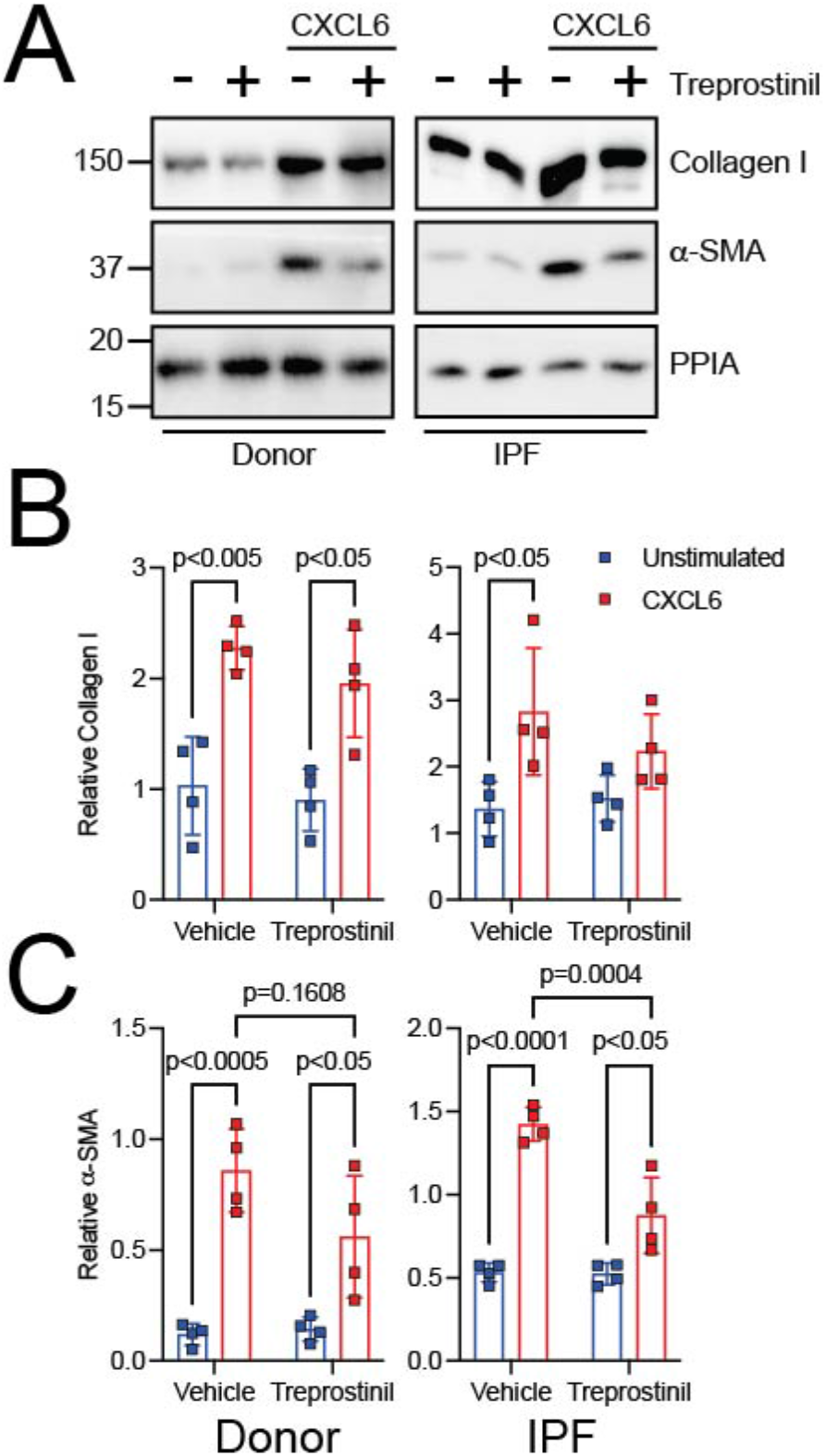
Treprostinil blocks CXCL6-induced increases in α-SMA but not collagen I. (A) Treprostinil can block CXCL6-induced increases in α-SMA but not collagen I Donor and IPF lung Fbs were stimulated with CXCL6 in the presence of the cAMP agonist treprostinil. IB was performed for Collagen I, α-SMA, and PPIA. Treprostinil blocked CXCL6-induced effects α-SMA but not collagen in both IPF and Donor Fbs. Representative blot of experiments with 4 independent IPF/Donor Fbs. (B) Densitometry of Collagen I from Donor (*left*) or IPF (*right*) fibroblasts normalized to PPIA from (*A*) expressed as mean ± SD. Post-hoc comparisons are noted in the figure. Data were analyzed by two-way ANOVA, followed by Fisher’s LSD post-hoc test. (C) Densitometry of α-SMA from Donor (*left*) or IPF (*right*) fibroblasts normalized to PPIA from (*A*) expressed as mean ± SD. Post-hoc comparisons are noted in the figure. Data were analyzed by two-way ANOVA, followed by Fisher’s LSD post-hoc test

### High levels of CXCL6 is associated with markers of more severe outcomes in IPF

Our data support that C*XCL6* gene expression is increased in IPF. We have also shown that CXCL6 promotes collagen synthesis in mice and in lung fibroblasts and supports myofibroblast differentiation as measured by expression of α-smooth muscle actin. If CXCL6 drives these pathologic effects in cells and in experimental animals, we reasoned that elevated levels of CXCL6 might drive the same effects in patients with IPF. And if so, CXCL6 levels would be associated with markers of poor outcomes in IPF. To explore this question, we returned to the LGRC data and considered only IPF patients. To determine if there is a correlation between *CXCL6* gene expression and pulmonary function, we plotted pulmonary function as a function of equal quartiles of *CXCL6* gene expression **(Figure 9A and Table 1)**. With increasing CXCL6 expression, both FVC (N=32-34 per quartile, for linear trend, *p*<0.0001) and DLCO (N=28-32 per quartile, *p*<0.0001) decreased. This suggests that increased CXCL6 gene expression is associated with worse pulmonary function in IPF. Because *CXCL6* gene expression is associated with more impaired lung function, we next determined if CXCL6, as a secreted protein can be measured at the protein level in patients. We found that plasma levels of CXCL6 were statistically higher in IPF patients (n=74) than in Donor controls (N=38), but the median difference between these groups was only 5 pg/mL. This difference was not considered biologically informative **(Figure 9B and Table 2)**. Because we observed CXCL6 expression in lung epithelial cells in IPF lungs, we reasoned that we might observe the highest levels of this protein in the bronchoalveolar lavage (BAL). Previously, it was shown that BAL from a small number of IPF patients showed elevated levels of CXCL6 compared to healthy volunteers (39). To answer this question, we quantified CXCL6 protein levels in the bronchoalveolar lavage (BAL) of patients with IPF. In BAL, the median CXCL6 concentration was 91.5 pg/mL in unaffected controls compared to 328 pg/mL in IPF patients (N=18 healthy controls and N=131 IPF patients, *p*=0.0001) (**Figure 9C; Tables 3 & 4**). Since the levels of CXCL6 were elevated 3.6-fold in IPF, we next determined if CXCL6 levels were associated with mortality risk (**Figure 9D-E**). Considering IPF patients only (Cohort 2018; N=120), employing Cox survival analysis, patients with above median levels of CXCL6, were at significantly greater risk for death or progression **(Figure 9D-E & Table 5)** than patients with below median levels. This risk persisted after controlling for age, sex, and FVC. To validate these findings, we determined the association of CXCL6 with death or progression in a second cohort. In 59 patients with IPF (Cohort 2021), we again observed that above-median levels of CXCL6 were associated with increased risk of death or progression compared to patients with below-median levels of CXCL6 **(Figure 9F-G)**. This difference also persisted after controlling for age, sex, and FVC (**Figure 9F-G & Table 5**). These data suggest that CXCL6 at the mRNA and protein levels are associated with markers of more severe outcomes in IPF.

**FIGURE 9.**
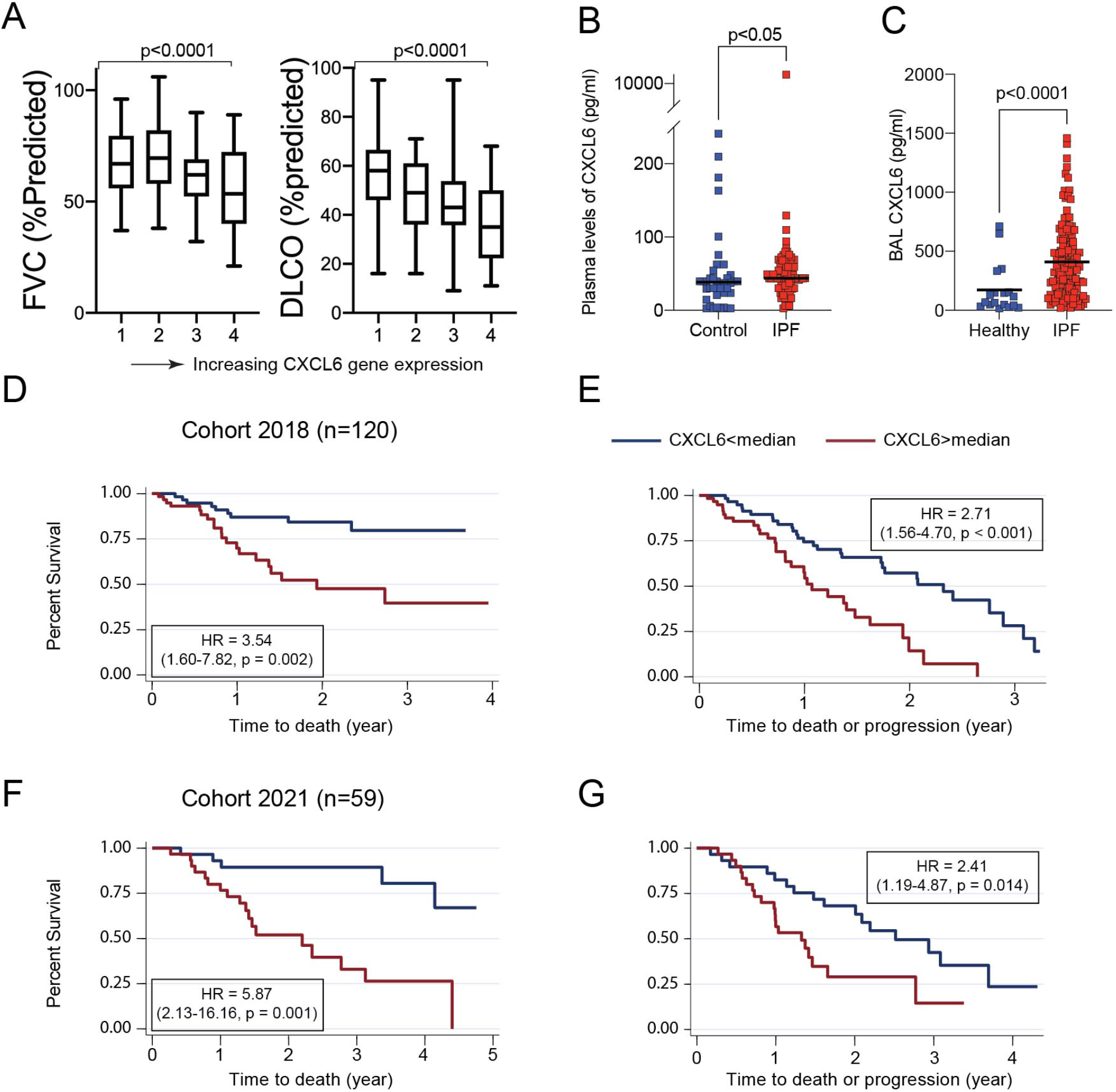
High levels of CXCL6 correlate with markers of worse outcomes in IPF. (A) Box and whisker plots of forced vital capacity (FVC) and diffusing capacity for carbon monoxide (DLCO) vs. assignment of IPF patients from the LGRC cohort into equal quartiles of CXCL6 gene expression. for FVC (left, N=32-34 per quartile, P for linear trend<0.0001) and DLCO (right, N=28-32 per quartile, P<0.0001). (B) CXCL6 levels are slightly elevated in the plasma of IPF patients (N=74) compared to normal controls (N=38) (P<0.05, Mann-Whitney) (C) CXCL6 levels are significantly elevated in the BAL of IPF patients (N=131) compared to normal controls (N=18) (P<0.0001, Mann-Whitney). (D) In the 2018 cohort, CXCL6 values exceeding the median value are associated with an increased risk of death (N=120, Hazard ratio (HR)=3.54 (1.60-7.82, P=0.002) and (E) Death or progression (HR=2.71 (1.56-4.70, P<0.001). (F) In the 2021 cohort, CXCL6 values exceeding the median value are associated with an increased risk of death (N=59, HR=5.87 (2.13-16.16, P=0.001) and (G) Death or progression (HR=2.41 (1.19-4.87, P=0.014)).

**TABLE 1.** Demographic and clinical characteristics of the LTRC Cohort

|  | **Control** | **IPF** | **p-value**<br>**(IPF v Control)** |
| --- | --- | --- | --- |
| N | 108 | 134 |  |
| Men (%) | 49 (45) | 94 (70) | <0.0001 |
| Women (%) | 59 (55) | 40 (30) |  |
| Age (SD) | 63.6<br>(11.4) | 56.4<br>(11.4) | <0.0001 |
| Ethnicity (%) |  |  |  |
| White | 100 (92) | 122 (91) | 0.95 |
| African American | 3 (2.8) | 5 (3.7) |  |
| Hispanic | 1 (0.9) | 1 (0.7) |  |
| Asian/Pacific Islander | 3 (2.8) | 2 (1.4) |  |
| Other | 1 (0.9) | 1 (0.7) |  |
| Smoking Status (%) |  |  |  |
| Never | 32 (30) | 47 (35) | 0.3 |
| Current | 2 (1.9) | 2 (1.5) |  |
| Ever | 63 (58) | 81 (60) |  |
| Pulmonary Function |  |  |  |
| FEV1, %predicted (SD) | 95 (12.6) | 70.5<br>(18.2) | <0.0001 |
| FVC, %predicted (SD) | 94.4<br>(13.1) | 63.4<br>(16.9) | <0.0001 |
| DLCO, %predicted (SD) | 84.1<br>(16.7) | 46.7<br>(19.2) | <0.0001 |

**TABLE 2.** Demographic and clinical characteristics of the Pitt cohort

|  | **Control** | **IPF** |
| --- | --- | --- |
| N | 38 | 74 |
| Age (year) | 50.4 (41-72) | 69.7 (49-87) |
| Sex (%) |  |  |
| Male | 22 (57.9) | 44 (59.5) |
| Female | 16 (42.1) | 26 (35.1) |
| Unknown |  | 4 (5.4) |
| Race (%) |  |  |
| White | 32 (84.2) | 67 (90.5) |
| African American | 2 (5.3) | 1 (1.4) |
| Other | 3 (7.9) |  |
| Unknown | 1 (2.6) | 6 (8.1) |
| Smoking (%) |  |  |
| Never | 23 (60.5) | 18 (24.3) |
| Current | 4 (10.5) | 3 (4.1) |
| Former | 9 (23.7) | 47 (63.5) |
| Unknown | 2 (5.3) | 6 (8.1) |
| Pulmonary Function* |  |  |
| FVC, %predicted (SD) |  | 71 (18.7) |
| DLCO, %predicted (SD) |  | 48 (14.3) |

\* based on 62 subjects

**TABLE 3.** Demographic and clinical characteristics of normal BAL controls (Hannover)

| Normal Controls |  |
| --- | --- |
|  | Cohort 2018<br>N = 18 |
| Age, median (IQR) year | 65 (61-73) |
| Male, n (%) | 11 (61) |
| FEV1%, median (IQR) | 106 (96-124) |
| FVC%, median (IQR) | 116 (104-132) |
| FEV1/FVC, median (IQR) | 73.9 (72.4-78.2) |
| CXCL6, median (IQR) | 91.5 (38-197) |

**TABLE 4.** Demographic and clinical characteristics of IPF BAL cohorts (Hannover)

| Patient characteristics |  |  |
| --- | --- | --- |
|  | **Cohort 2018**<br>N = 120 | **Cohort 2021**<br>N = 59 |
| Age, median (IQR) year | 74 (65-78) | 71 (62-79) |
| Male, n (%) | 96 (82) | 45 (76) |
| Current Smoker, n (%) |  | 2 (4) |
| Baseline FVC%, median (IQR) | 69 (58-83) | 71 (60-89) |
| Baseline DLCO%, median (IQR) | 52 (39-62) | 54 (41-70) |
| CXCL6, median (IQR) | 324 (143-561) | 1168 (807-2045) |
| Follow up time, median (IQR)<br>year | 1.2 (0.6-2.2) | 2.0 (1.3-3.3) |
| Death, n (%) | 29 (24)* | 23 (39) |
| Progressed, n (%) | 31 (26) | 24 (41) |
| * Including two patients who received lung Tx |  |  |

**TABLE 5.** Survival analysis of IPF BAL cohorts

| Survival analysis |  |  |
| --- | --- | --- |
|  | Cohort 2018 | Cohort 2021 |
| **Death** |  |  |
| Unadjusted HR (95%CI) | 3.54 (1.60-7.82, P = 0.002) | 5.87 (2.13-16.16, P = 0.001) |
| Adjusted HR (95%CI)* | 2.82 (1.26-6.28, P = 0.011) | 4.47 (1.52-13.10, P = 0.006) |
| **Death or progression** |  |  |
| Unadjusted HR (95%CI) | 2.71 (1.56-4.70, P < 0.001) | 2.41 (1.19-4.87, P = 0.014) |
| Adjusted HR (95%CI)* | 2.78 (1.55-4.90, P = 0.001) | 2.18 (1.02-4.62, P = 0.043) |
| * For age, sex, **and** FVC |  |  |

## DISCUSSION

In this study, we have shown that CXCL6—a chemokine which signals via Gαi—drives fibrotic effector functions and correlates with markers of poor clinical outcomes in IPF. These data suggest that CXCL6 is not only a potential biomarker of IPF but also an important therapeutic target. We have shown that CXCL6 is predominately expressed in lung epithelial cells and may signal in fibroblasts and in inflammatory cells. In animals, administration of mCXCL5 (the closest match to human CXCL6) increased collagen synthetic activity in the lung—in the presence and absence of bleomycin. The increased collagen synthetic activity was accompanied by a significant accumulation of neutrophils. *In vitro*, we found that CXCL6 stimulation increased cellular levels of collagen I and α-SMA. These effects could be blocked by silencing of the CXCL6 receptors CXCR1 and CXCR2 or by chemical inhibition with the small molecule reparixin. CXCL6-induced expression of α-SMA, but not collagen, was blocked by the Gαs/cAMP agonist treprostinil suggesting that CXCL6-induced collagen was independent of cAMP. And, in two cohorts of IPF patients, we found that above-median levels of CXCL6 were associated with significantly increased risk of death or progression. These data are supportive of a previous study with a limited number of participants showing that CXCL6 levels are increased in the BAL of IPF patients (39).

Our findings may suggest a rationale for testing CXCR1/2 blockade as an anti-fibrotic strategy in IPF. This does, however, raise the question on the role of the neutrophil in pulmonary fibrosis. The best-known function of the CXCL6 family is to drive neutrophil chemotaxis (40). If CXCL6 is a significant mediator of pulmonary fibrosis, then this would suggest that neutrophils play a critical role in disease. In experimental lung injury, some have observed that neutrophils are necessary, while others have observed that they are dispensable (41–44). Blockade of CXCR2 with DF2162 blocked bleomycin-induced pulmonary fibrosis (45). Neutrophils, as sources of proteases (46), have been associated with myofibroblast differentiation (47) and bleomycin-induced fibrosis (43, 48). The loss of neutrophils may potentiate pulmonary fibrosis (41). Yet, neutrophils are also necessary for experimental hypersensitivity pneumonitis (42). The association of the IL-8 with IPF prognosis (49) and the association of neutrophils in the BAL with early mortality (50, 51) are suggestive of a critical role for neutrophils in driving IPF. An intriguing idea suggests that turnover of collagen, mediated by neutrophil-derived proteases, can generate acetylated proline-glycine-proline fragments which have been shown to bind to CXCR2 which may further recruit neutrophils to the site of injury (52). In support of a direct effect on the mesenchymal compartment, we have shown that CXCL6 signaling directly impacts fibroblast function. Our data suggest that further studies are needed to understand the role of neutrophils in promoting fibrotic injury.

How do we place these findings in context? CXCL6 is a member of the IL-8 family. IL-8 has previously been shown to be a marker of mortality risk in IPF (49). Increased levels of another family member, CXCL1, have been associated with IPF exacerbations (53). IL-8 has been shown to increase proliferation and motility in a community of IPF mesenchymal progenitor cells (54). Multiple members of the IL-8 family are up-regulated in both *in vitro* and *in vivo* models of telomere dysfunction in epithelial cells (55), which serves as a potential link between dysregulated CXCL6 and the pathogenesis of familial and sporadic pulmonary fibrosis (56, 57). We observed that elevated levels of CXCL6 is associated with low expression of *RXFP1.* Thus up-regulation of CXCL6 family is coupled to down-regulation of several Gαs-coupled receptors, which may represent a critical shift in signaling away from “anti-fibrotic” Gαs signaling to “pro-fibrotic” Gαi signaling (10, 58). Gαi-coupled GPCRs are upregulated globally in IPF lungs while expression of Gαs-coupled GPCRs is relatively lower (22). Gαs-coupled receptors (that elevate cAMP tone) such as *PTGIR* (Prostacyclin I2 receptor, IP2), *PTGER2* (Prostaglandin E receptor 2, EP2) (59) and *RXFP1* (9) showed low expression in IPF fibroblasts compared to donor fibroblasts. Gαi-coupled receptors, such as *LPAR1* (*LPA_1_*) and *LPAR2* (Lysophosphatidic acid receptors 1 and 2) (21, 61), that decrease cAMP tone, are highly expressed in IPF compared to donors. In the context of these studies, we suggest that dysregulation of CXCL6 is a critical element of a synchronized wave of CXCR1/2 signaling which originates from diverse cellular sources to promote matrix synthesis and to stimulate contractility. While we have explored expression of collagen and α-smooth muscle actin as markers of the pro-fibrotic functions of CXCL6, signaling via CXCR2 in fibroblasts can induce a senescent phenotype (62) and has been shown to upregulate the profibrotic cytokine CTGF (63).

Our studies do leave several mechanistic questions unanswered. Recently, the cAMP agonist treprostinil, in a phase 2 study of patients with interstitial lung disease (ILD) and pulmonary hypertension, was associated with improved exercise based on 6-minute walk test when compared to placebo (64). We have observed that CXCL6-induced collagen is independent of treprostinil and by extension, independent of cAMP. Therefore, the other arm of G protein signaling, via G*βγ* subunits, may also promote fibrotic signaling independent of cAMP. G*βγ* signaling may lead to increases in intracellular and activation of phospholipase C (PLC)-β (65). Activation of PI3K*γ*, which is activated upon dissociation of the G*βγ* subunits (66), has been shown to complex with and promote TRPV4 translocation to the cell membrane where it functions as a Ca^2+^ channel to drive up intracellular calcium in response to mechanosensitive stimuli (67, 68). Further studies are needed to explicate the parallel signaling pathways initiated by CXCL6 on fibrotic effector functions in IPF.

Finally, can we employ CXCL6 as a biomarker? We did not detect differential levels CXCL6 in the plasma of patients with IPF compared to donor controls. Plasma would be the easiest body fluid to sample for biomarkers. One interpretation to explain the difference between the BAL and the plasma is that the kinetics of CXCL6 in the peripheral blood remains unknown. It is notable that the observed levels of IL-8 in IPF are similar to the CXCL6 levels we measured in the plasma (49). We did, however, detect elevated levels of CXCL6 in the bronchoalveolar lavage. Translating CXCL6 in the BAL into a clinical biomarker would be challenging as this is a rare practice for the diagnosis and management of IPF in United States. Nevertheless, BAL is still recommended by the most recent consensus guidelines (69). The performance of BAL for detection of biomarkers is likely dependent on technique and the operator. Recent guidelines suggest that bronchoscopy should adopt uniform standards of practice (70). Our study supports the role of well-performed BAL analysis as a highly informative resource for understanding both the clinical behavior and mechanisms of interstitial lung disease (53, 71, 72).

In summary, we have shown that elevated levels of CXCL6 are associated with markers of poor clinical outcomes in IPF. This signal also drives fibrotic pathology *in vivo* and in cultured human fibroblasts *in vitro*. We suggest that targeting CXCR1/2 family signaling may be a potentially important therapeutic strategy in IPF.

## EXPERIMENTAL PROCEDURES

### Ethical review

Collection of human lung tissue was approved by the Institutional Review Board and the Committee for Oversight of Research and Clinical Training Involved Decedents of the University of Pittsburgh. This study was conducted in accordance with University of Pittsburgh IRB protocols 20070385, 19040326, 20030050, 20030223, 14120072, 0411036, CORID protocol 746, and IACUC 18103979. Informed written consent to the internal ethics review board-approved clinical study protocol (Hannover Medical School IRB protocol # 2923-2015) was obtained from each individual before participation in this study, in accordance with the Declaration of Helsinki.

### Isolation of human lung tissue

Human lung tissues were obtained from excess pathologic tissue after lung transplantation and organ donation, under a protocol approved by the University of Pittsburgh Institutional Review Board as previously described(73). Human lung tissue pieces were cut from the upper and lower lobes and placed in 2% paraformaldehyde (PFA) for fixation. Sections were then embedded in paraffin for sectioning.

### Primary cell culture

Donor human fibroblasts were isolated from lungs that appeared to have no injury by histology but were deemed unacceptable for lung transplantation. IPF lung fibroblasts were obtained from patients either at explant or at autopsy(^74^). Briefly, lung tissue from explanted IPF lungs and age-sex matched normal donors was collected from the lower lobes to find age and sex of the lines included in the experiments (Mean age for IPF 65y ± 5 and 64y ± 3 for controls p=0.60). Enzymatic digestion (0.05% trypsin; GIBCO) was used to isolate human lung fibroblasts. Fibroblasts were grown with Dulbecco’s modified Eagle’s medium supplemented with 10% fetal bovine serum (Corning) containing 10% of fetal bovine serum (Atlanta Biologicals) and 1% of antimycotic-antibiotic (GIBCO). Cells were cultured at 37°C, 5% CO_2_ and then expanded to select a homogeneous fibroblast population (experiments were performed with lines between passages 5 and 9).

### Microarray analysis

A full description of the RNA isolation and microarray procedures have been previously described(75, 76). We used publicly-available data from the Lung Genomics Research Consortium (LGRC) (12). 134 IPF patients and 108 normal controls were selected. These data and methods are also available on the Gene Expression Omnibus Database (GEO) under reference number GSE47460. The data were normalized using a cyclic loess algorithm from the Bioconductor suite of R tools (77). False discovery rate (FDR) was calculated according to the method of Benjamini and Hochberg (78).

### cAMP assay

Donor and IPF lung fibroblasts (N>6) were plated at 5K cell/well overnight. Where indicated, cells were incubated with recombinant human CXCL6 (R&D) ranging from 1 nM to 1 μM. Between 16h and 24 h, cells were processed to assess cAMP levels using the luminescent cAMP Glo assay (Promega). For dose–response curves, Fibroblasts were incubated with a range of CXCL6 concentrations from 1 nM to 1 μM. Subsequently, cells were lysed and the protocol was followed according to the manufacturer’s recommendation (Promega). Luminescence was measured using a microplate luminometer (Biotek Synergy H1 plate reader). The cAMP levels (RLU; relative luminescence unit) were evaluated by subtracting the CXCL6 conditions from the untreated conditions. Luminescence was correlated to the cAMP concentrations by using a cAMP standard curve (0-5 μM). Each experiment had 3 technical repeats per condition and was independently repeated at least three times. To obtain IC_50_, dose–response data were fit to a sigmoidal dose–response equation using nonlinear least-squares regression on Graphpad Prism software.

### Bleomycin-induced lung injury and treatment with mCXCL5

Eight-week-old female C57BL/6 mice with body weight of 20-25g were purchased from the Jackson Laboratory (Bar Harbor, ME). The mice were anesthetized with isoflurane in an anesthesia chamber. Bleomycin (Sigma-Aldrich, St. Louis, MO) was administered at 2 U/kg by inhalation on day 0 (79). After bleomycin injury, Recombinant mouse CXCL5/LIX (R&D) or the vehicle (sterile water) was administered by transnasal aspiration at 0.15 mg/kg (∼3 μg) on days 7, 10, and 13 after injury. Mice were sacrificed on day 14. Details of flow cytometric analysis of the bronchoalveolar lavage have been previously published (9, 18).

### Flow Cytometry

Mouse BAL cells were counted by trypan-blue exclusion method followed by incubation with CD16/CD32 antibody (BD Pharmingen) for 30 min on ice. Cells were then stained with the following: Ly6G-FITC (BD Pharmingen; clone IA8), CD68-PE (Bio Legend; clone FA-11), CD3-APC (BD Pharmingen; clone 145-2011), B220-PeCy7 (BD Pharmingen; clone RA3-6B2). Samples were acquired on a FACSAria II or Fortessa (BD Biosciences, San Jose CA) and analyzed by *FlowJo* software (Tree Star, Ashland OR).

### Histology

The right lungs were inflated and fixed with 4% paraformaldehyde (USB Corporation, Cleveland, OH) at 25 cm H_2_O pressure. Lungs were then excised and fixed by immersion in 4% paraformaldehyde for at least 24h before paraffin embedding and sectioning. Immunohistochemistry was performed on formalin-fixed paraffin embedded lungs following Tris-EDTA pH 9.0 antigen retrieval with an antibody that recognizes CXCL5/6 (Abcam). Routine hematoxylin and eosin (H&E) and Masson’s trichrome staining was performed by the University of Pittsburgh histology service.

### Single cell RNASeq analysis

Previously reported UMI count data of scRNAseq was downloaded from the GEO accession GSE136831. Analysis was performed in R version 3.6.1 using the Seurat package 3.0.4. Cells from COPD samples or previously identified as multiplets were discarded. UMI counts were scaled to 10,000 UMIs per cell, then log transformed with a pseudocount of 1. The top 1,500 genes were selected based on variance using the Seurat implementation *FindVariableFeatures*. These genes were then centered and scaled using Seurat’s implementation *ScaleData* and total UMI and percent mitochondrial genes as variables to regress out; scaled data was subsequently used for principal component analysis. We then selected the top 50 principal components for visualization – however we avoided principal components 29 and 30 because they represented subject outlier signals. We constructed a two-dimensional uniform manifold approximation and projection (UMAP) with Seurat’s *RunUMAP* implementation under the following altered parameters: 50 nearest neighbors, local connectivity: 5, minimum distance: 0.4, spread: 1.5, repulsion strength: 1.5, 1000 epochs and the seed 7. All cell type assignments are those reported in the original analysis. Density plots were made with the R packages *ggplot2* and *ggridges*. Statistical comparisons of CXCL6 were performed using wilcoxon rank-sum tests on normalized data with Seurat’s implementation *FindMarkers* for cell-cell comparisons, or R’s base *wilcox.test* implementation on average per subject normalized expression for sample-sample comparisons.

A second single cell RNA sequencing dataset of three control lungs, donor lungs not suitable for lung transplant, and three paired upper and lower explanted IPF lungs (GEO Accession: GSE128033) was used for analyzing cell type-specific *CXCL6* expression using the exact method described (15). Lung tissue collection, single cell isolation, sequencing, data cleaning, normalization, clustering, and cluster identification was described in detail in the original study (15). Violin plots were generated by comparing the normalized single cell data between the normal controls and the lower IPF lobes for each cluster. Each dot represents the level of gene expression for a particular cell with the group’s expression distribution visualized as the violin. Violin plots were generated using the Seurat package in R. The details have been published previously (80).

### Sircol assay for acid-soluble collagen

Mouse lungs were perfused through the right ventricle with PBS. The left lungs were excised and flash-frozen in liquid nitrogen, followed by lyophilization in preparation for collagen determination by the Sircol collagen assay (Biocolor life science, Belfast, UK). Lungs were homogenized in 0.5 M acetic acid with protease inhibitors (Sigma-Aldrich). The homogenate was pelleted, and the supernatant was run across a QIAshredder column (Qiagen). The lung–acetic acid mixture was incubated with Sircol Red reagent. The collagen-dye complex was pelleted, and the precipitated material was redissolved in NaOH. O.D_540_ was recorded using a microplate reader.

### Transfection of siRNA

ON-TARGETplus smartpool siRNAs for CXCR1/2, and negative control scramble siRNA were purchased from Dharmacon. Fibroblasts were transfected with siRNA using HiperFect transfection reagent (Qiagen). Briefly, transfection complexes were generated by mixing 0-10 nM of siRNA mimic/ control mimic according to the manufacturer’s protocol. After 10-minute incubation at room temperature, the transfection complexes were added drop-wise onto 2 × 10^5^ cells per well of a 6-well plate in 2000 μL of culture medium with 1% FBS but without antibiotics. Cells were transfected on the day of plating (Day 0) and again on Day 1. Brefeldin A was used at a concentration of 10 μg/ml for 30 min before harvesting the cells.

### Western Blotting

CXCL6 protein expression was analysed using Western blot. Western blotting was performed as described previously(9). Anti-CXCL5/6 antibody (ab198505; Abcam), Mouse anti-β-actin (sc-47778; Santa Cruz Biotechnology), anti-collagen I antibody (PA5-29569; Invitrogen) and anti-cyclophilin (AB178397; Abcam) were used. Chemiluminescence reagent was purchased from Advansta (San Jose, CA). Densitometry analyses were performed using the *imageJ* software.

### Analysis of human bronchoalveolar lavage

Patients with presumed IPF at Hannover Medical School underwent bronchoscopy and bronchoalveolar lavage (BAL) during the routine diagnostic work-up. The retrieved BAL was immediately processed in a closed laboratory. The fluid was pooled, filtered through two layers of gauze, and centrifuged at 500 g for 10 minutes at 4°C. The cells were counted and cell smears were stained with May-Grunwald-Giemsa stain (Merck, Darmstadt, Germany)(71, 81). BAL was subjected to multiplex Luminex analysis (Bio-Plex 200, BioRad).

### Statistical Analysis

All data are shown as mean ± S.D. Statistical significance was assessed by performing resampling boot-strapped test, Mann-Whitney test, one-way and two-way analysis of variance (ANOVA), where appropriate, followed by the Fisher’s LSD post-hoc analysis for multiple comparisons in GraphPad Prism 9.0 (San Diego, CA) and Stata 16.0 (StataCorp, College Station, TX) unless otherwise stated. Statistical analysis is indicated in the figure legends.

## Supporting information

Summary statistics for %Cells w/ expression

Results of Wilcoxon rank-sum tests for differential

Results of CXCL6 spearman correlation

## Acknowledgements

The authors thank Justin A. Dutta for technical assistance.

## Conflict of interest

NK served as a consultant to Biogen Idec, Boehringer Ingelheim, Third Rock, Pliant, Samumed, NuMedii, Theravance, LifeMax, Three Lake Partners, Optikira, Astra Zeneca, Veracyte, Augmanity and CSL Behring, over the last 3 years, reports Equity in Pliant and a grant from Veracyte, Boehringer Ingelheim, BMS and non-financial support from MiRagen and Astra Zeneca. NK has IP on novel biomarkers and therapeutics in IPF licensed to Biotech. Other authors declare no conflict of interest. Dr. Lafyatis reports grants from Bristol Meyer Squib, Corbus, Formation, Moderna, Regeneron, Pfizer, and Kiniksa, outside the submitted work; and served as a consultant with Bristol Meyers Squibb, Formation, Sanofi, Boehringer-Ingelheim, Merck, and Genentech/Roche. Dr. Prasse reports personal fees and non-financial support from Boehringer Ingelheim, personal fees and non-financial support from Roche, personal fees from Novartis, personal fees from AstraZeneca, personal fees from Amgen, personal fees from Pliant, personal fees from Nitto Denko, outside the submitted work.

## Author Contributions

JT, HB, DS, JA, TT, RL, PSB, CJ, NK, XL, and YZ conducted the *in vitro* and *in vivo* experiments and analysed the results. HB, JT, TT, RL, PB, CK, MN, AP, and DJK analyzed the results. JS, MR, XL, TC, and YZ provided tissue and cells for analysis. Data from clinical samples were analyzed by NK, YZ, RMA, BS, MN, AP, and DJK. AP, DJK, JT, and HB conceived and planned experiments. HB, JT, AP, and DJK wrote the manuscript.

## FOOTNOTES

This work was supported in whole or part by R01 HL 126990, P50 AR 060780-06A1, the Violet Rippy 5K for Pulmonary Fibrosis Research Fund, the Massaro Family Pulmonary Fibrosis Research Fund, and the Simmons Genetics Research Fund. NIH NHLBI grant 5T32HL007563-32 to HB. NIH NHLBI grants R01HL127349, R01HL141852, U01HL145567, UH2HL123886 to NK, and a generous gift from Three Lakes Partners to NK. AP is supported by KFO311, and E-RARE IPF-AE. 2P50 AR060780 to RL.

^§^ These authors contributed equally to this work.

The abbreviations used are: CXCL6, C-X-C Motif Chemokine Ligand 6; RXFP1, relaxin/insulin-like family peptide receptor 1; TGF, transforming growth factor; DMEM, Dulbecco’s modified Eagle’s medium; GPCR, G-protein coupled receptor; ANOVA, analysis of variance; IPF, Idiopathic pulmonary fibrosis

