## Supplemental material for "High CXCL6 drives matrix expression and correlate with markers of poor outcome in IPF"

^1^From the Dorothy P. and Richard P. Simmons Center for Interstitial Lung Disease, Division of Pulmonary, Allergy and Critical Care Medicine, University of Pittsburgh School of Medicine, Pittsburgh, PA-15213; ^2^Fraunhofer ITEM, Nicolai-Fuchs-Strabe 1, 30625 Hannover, Germany & DZL BREATH; ^4^Pulmonary, Critical Care and Sleep Medicine, Ohio State University, Columbus, OH; ^3^Department of Respiratory Medicine, Hannover Medical School, Carl-Neuberg Strasse 1, 30625 Hannover, Germany & German Centre for Lung Research (DZL), Biomedical Research in End-stage and Obstructive Lung Disease Hannover; ^4^Pulmonary, Critical Care and Sleep Medicine, College of Medicine, Ohio State University; ^5^Division of Rheumatology and Clinical Immunology, University of Pittsburgh, Pittsburgh, PA, USA; ^6^Department of Medicine, University of Pittsburgh, Pittsburgh, PA, USA; ^7^Pulmonary, Critical Care and Sleep Medicine, St. Elizabeth medical center, Brighton, MA; ^8^Pulmonary, Critical Care and Sleep Medicine, Yale School of Medicine, New Haven, Connecticut; ^9^Unit for Interstitial Lung Diseases, Department of Respiratory Diseases, University Hospitals Leuven and Department of Chronic Diseases, Metabolism, and Ageing, KU Leuven, Leuven, Belgium, ^10^Starzl Transplantation Institute, University of Pittsburgh, Pittsburgh, PA; and ^11^Division of Pulmonary, Allergy and Critical Care Medicine, University of Pittsburgh School of Medicine, Pittsburgh, PA-15213.

*To whom all correspondence should be addressed: Simmons Center for Interstitial Lung Disease. Division of Pulmonary, Allergy, and Critical Care Medicine. University of Pittsburgh School of Medicine, 200 Lothrop St, Pittsburgh. Tel.: 412-624-7444; Fax: 412-624-1670; E-mail: [](http://)

**Keywords:** Biomarker, lung fibroblasts, IPF, CXCL6, Relaxin, pulmonary fibrosis

**Supplementary Figure S1:**

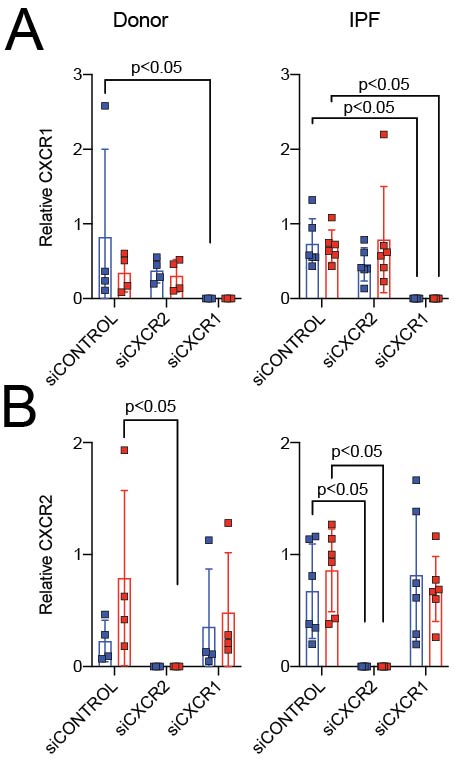

(A) Densitometry of CXCR1 from (*A*) data are expressed as mean ± SD., normalized to PPIA. Donor and IPF lung fibroblasts were incubated with CXCR1 or CXCR2 siRNA and stimulated with CXCL6. Transfection of siCXCR2 significantly decreased levels of CXCR2 protein compared to scramble control in both vehicle or CXCL6 treatment (Donor-p<0.05, N=4; IPF-p<0.05, N=6). Data were analyzed by two-way ANOVA, followed by Fisher’s LSD post-hoc test.
(B) Densitometry of CXCR2 from (*A*) data are expressed as mean ± SD., normalized to PPIA. Donor and IPF lung fibroblasts were incubated with CXCR1 or CXCR2 siRNA and stimulated with CXCL6. Transfection of siCXCR1 significantly decreased levels of CXCR1 protein compared to scramble control in both vehicle or CXCL6 treatment (Donor: p<0.05, N=4; IPF: p<0.05, N=6). Data were analyzed by two-way ANOVA, followed by Fisher’s LSD post-hoc test.

**Supplementary Figure S2 (uncut blots for Figure 1)**

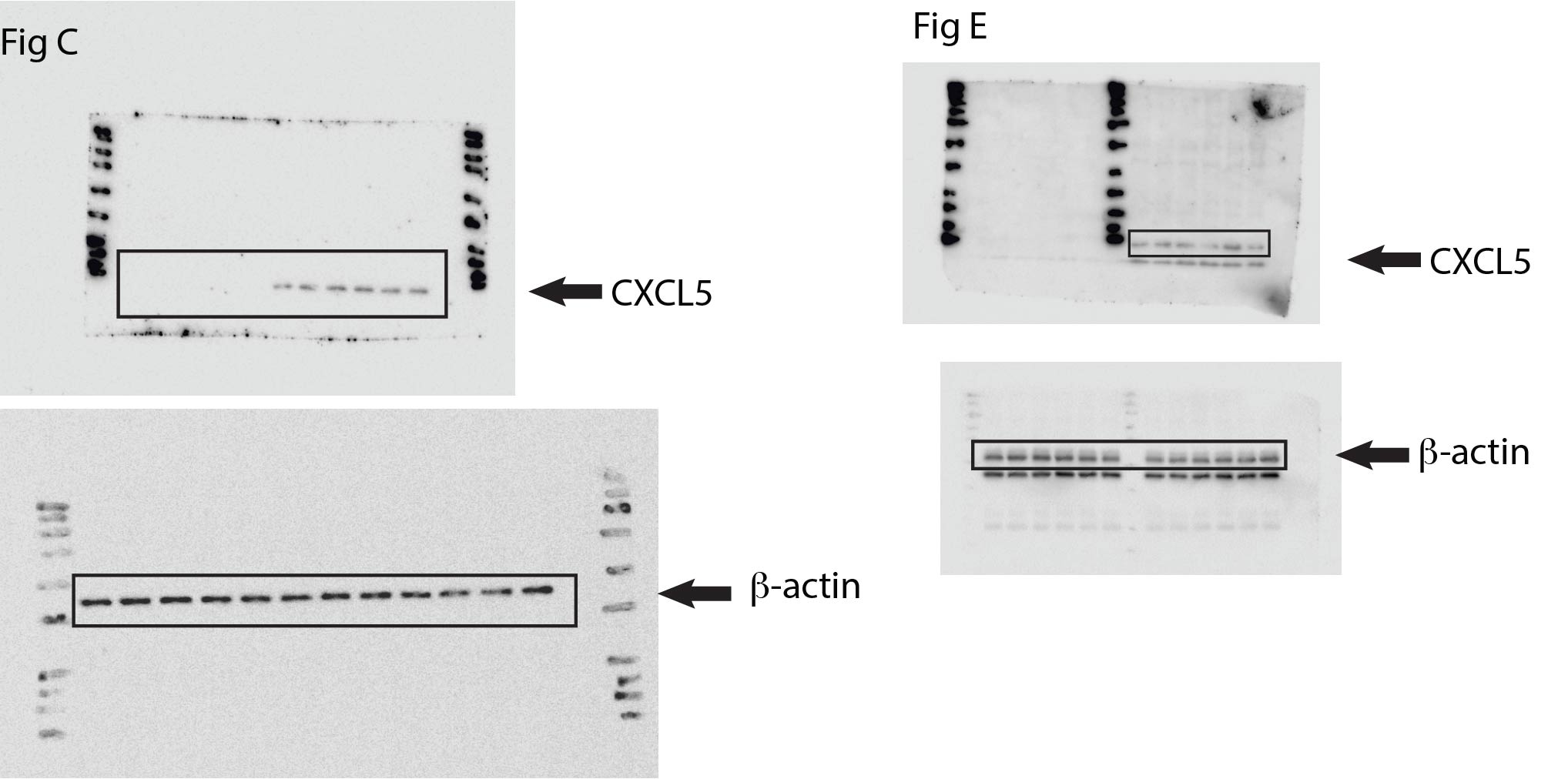

**Supplementary Figure S3 (uncut blots for Figure 5)**

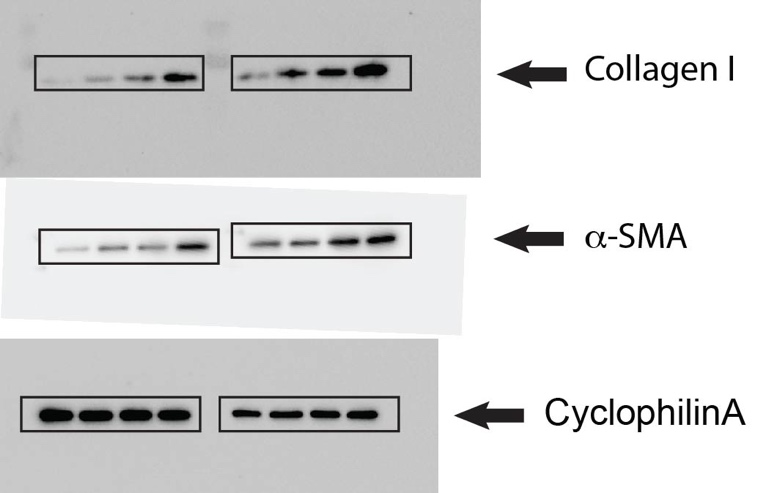

**Supplementary Figure S4 (uncut blots for Figure 6)**

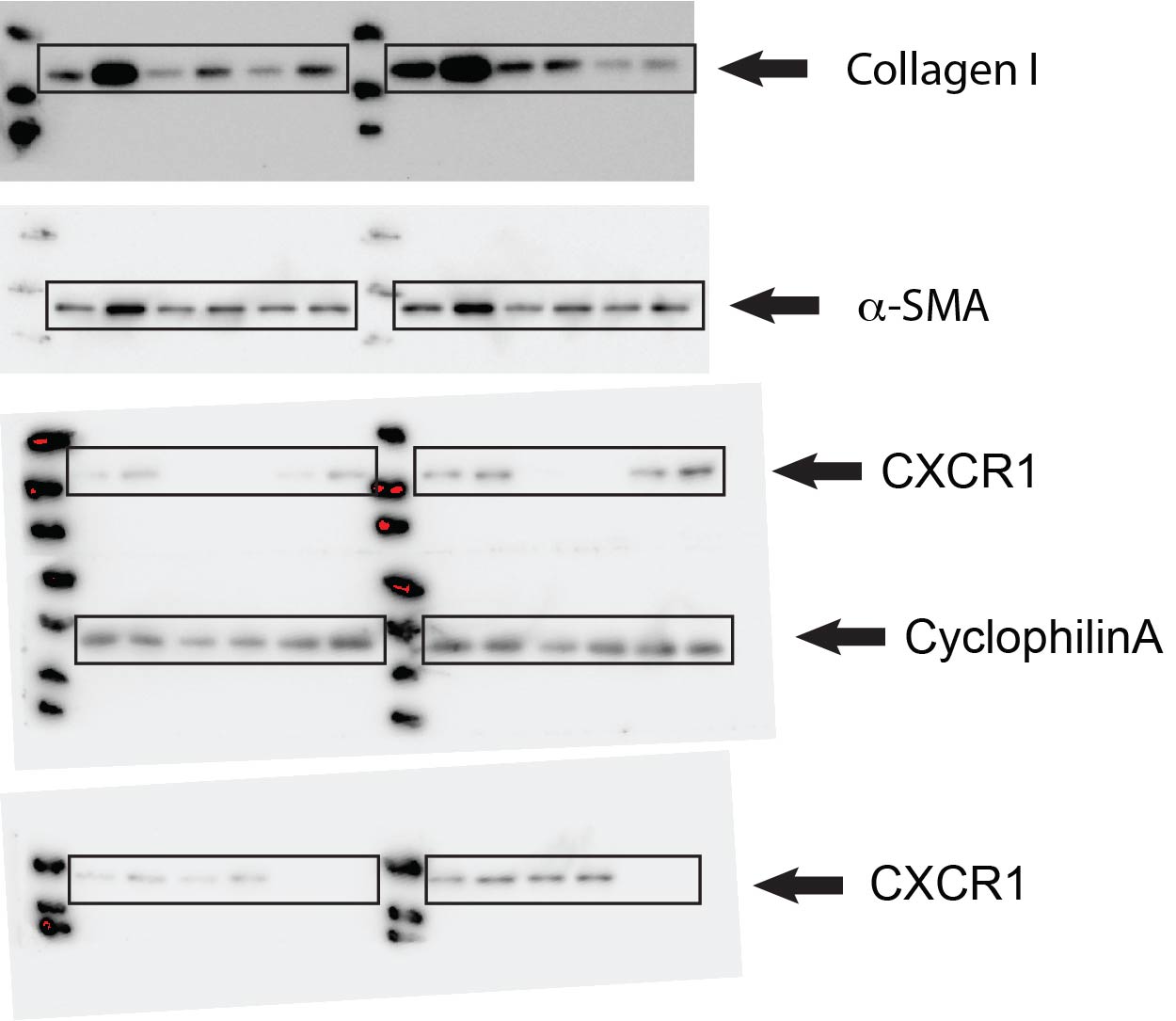

**Supplementary Figure S5 (uncut blots for Figure 7)**

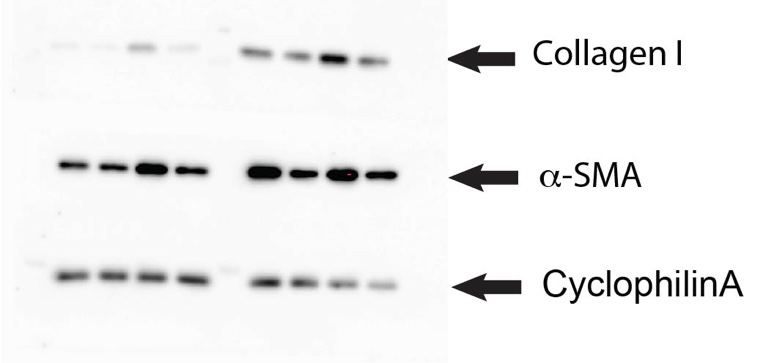

**Supplementary Figure S6 (uncut blots for Figure 8)**

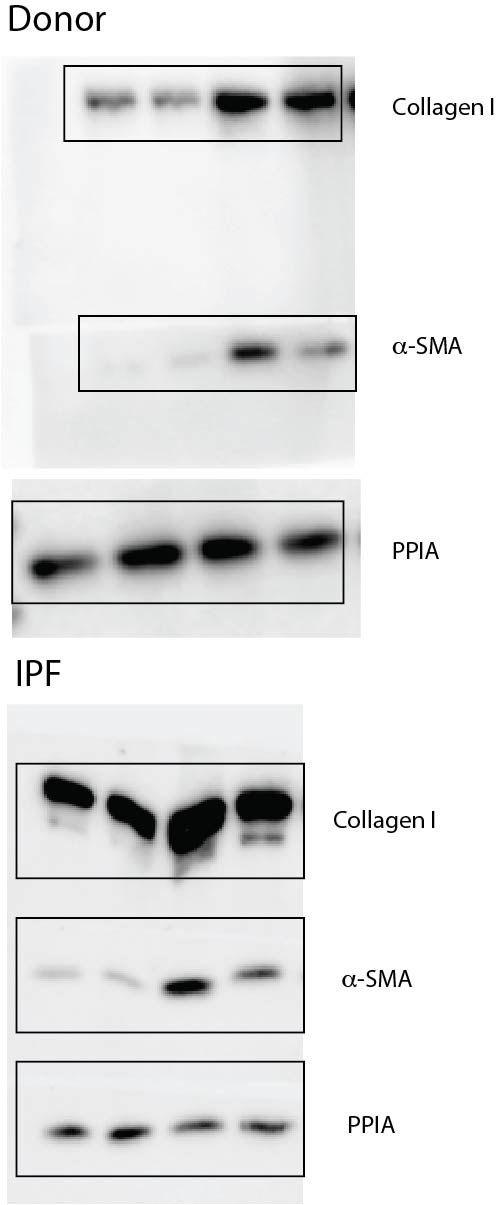

**Supplemental files**

**CXCL6_CellTypeExpressionSummaryStats.txt**

Summary statistics for %Cells w/ expression, average raw and normalized CXCL6 expression across all cell types, split by IPF or control.

**IPFvsCtrl_scRNAseq_CXCR6_WilcoxRankSum.txt**

Results of Wilcoxon rank-sum tests for differential expression at either the average/subject **or** cell level, for each of the top CXCL6-expressing cell-types, IPF versus Control.

**CXCL6_spearmanCorrelations_topCellTypesIPF.txt**

Results of CXCL6 spearman correlation tests of all genes expressed in > 5% of each cell type. Only IPF cells are used for this analysis, only the top 4 expressing cell types are evaluated (Goblet, Aberrant_Basaloid, Club and Fibroblast). Genes with fdr-adjusted theoretical p-value values < 0.05 were discarded.
